# Isolation and Purification-Free Digital Single-Small Extracellular Vesicle Biosensing with Scalable Plasmonic Arrays

**DOI:** 10.64898/2026.04.30.721846

**Authors:** Mohammad Sadman Mallick, Saswat Mohapatra, Abhay Kotnala, A B M Arafat Hossain, Wei-Chuan Shih

## Abstract

Recent advances in plasmonic biosensing and imaging have enabled label-free analysis of single biological nanoparticles. We previously developed PlAsmonic NanOapeRture lAbel-free iMAging (PANORAMA) for isolation and purification-free, digital counting and precise localization of antibody captured, tetraspanin-positive small extracellular vesicles (sEVs), with complementary fluorescence interrogation of surface and intravesicular biomarkers for quantitative molecular profiling. The fact that no isolation and purification is needed represents a crucial advantage because various specificity, efficiency, and time-consumption issues hinder quantitatively reproducible extraction of sEVs from biological fluids. PANORAMA achieves ultrahigh refractive-index sensitivity through arrayed gold nanodisks on invisible substrates (AGNIS) fabricated by nanosphere lithography (NSL). However, despite its simplicity and low cost, NSL is frequently constrained by poor large-area uniformity, which hinders scalable fabrication. Here, we introduce nanosphere settling lithography (NSSL) as an alternative to the gold-standard Langmuir–Blodgett trough (LBT) process, enabling highly uniform, large-area monolayers with reduced process stringency. AGNIS fabricated via NSSL exhibits high refractive-index sensitivity with low spatial variability across 60 mm × 24 mm substrates, sufficient for 60-well in standard 384-well plate format. The platform demonstrates exquisite sensitivity through PANORAMA digital counting and sizing of 25, 50, and 100 nm polystyrene beads, as well as single-vesicle characterization of antibody captured, tetraspanin-positive sEVs derived from H460 lung cancer cells. Combined PANORAMA and fluorescence imaging enables quantitative analysis of microRNA-21 (miR-21)-associated fluorescence among antibody captured sEV-like particles from liver cancer patient plasma, allowing label-free digital counting after selective affinity capture and simultaneous molecular profiling of the captured vesicle subset. Collectively, NSSL enables uniform, high-performance plasmonic biosensing over large areas, providing a scalable and economical pathway for high-throughput, digital analysis of antibody captured single sEVs and translational liquid biopsy applications.

---

Localized surface plasmon resonance (LSPR) sensors enable real-time, label-free detection of biological targets by transducing local refractive index changes into measurable spectral shifts arising from near-field–enhanced plasmonic nanostructures^1–4^. Periodic gold nanodisk arrays are particularly attractive for LSPR sensing due to their strong field confinement, tunable resonance, and compatibility with large-area fabrication^5–11^. When combined with advanced optical imaging modalities, such plasmonic platforms enable high-resolution analysis of nanoscale biological objects. PlAsmonic NanO-apeRture lAbel-Free iMAging (PANORAMA) is one such technique that leverages high-density arrayed gold nanodisks on invisible substrates (AGNIS) to achieve diffraction-limited, high-sensitivity imaging and digital counting of single nanoparticles^12^. It has been successfully applied to characterize small extracellular vesicles (sEVs), which are emerging biomarkers in cancer and other diseases. Digital detection of individual sEVs enables the identification of population heterogeneities that are otherwise masked in ensemble measurements. This approach holds significant promise for clinical diagnostics, as single vesicles carry molecular signatures reflective of their parent cells^13^. However, reliable characterization and detection of sEVs remain challenging due to their nanoscale dimensions and the inherent limitations of conventional analytical techniques. Although many techniques have been proposed to perform single sEV analysis, most of them require “clean” samples where sEVs are isolated or extracted in advance^14–17^. PANORAMA overcomes these barriers by enabling single-sEV counting, cluster analysis, and three-dimensional localization with ∼21 nm lateral and 7–9 nm axial accuracy^18^. In clinical studies, PANORAMA achieved 99.5% sensitivity and 97.3% specificity for cancer detection by directly counting individual sEVs in 20 μL human blood plasma without prior sEV extraction^19^.

Nanosphere lithography (NSL) offers a simple and cost-effective route to fabricate large-area plasmonic sensors without reliance on conventional lithography systems which can be quite costly ^20–23^. Despite these advantages, the performance of NSL-based plasmonic sensors is critically limited by the quality of the colloidal monolayer template. Achieving uniform, defect-free monolayers over inch-scale areas, particularly using submicron particles, remains a major challenge. Numerous NSL techniques have been developed to improve large-area ordering and reduce defects, including spin coating, dip coating, dry rubbing, convective self-assembly, and interfacial coating^24–27^. However, many of these techniques suffer from evaporation-induced defects, substrate dependence, poor reproducibility, or stringent process control, limiting their scalability and reliability for biosensing applications. Interfacial assembly using Langmuir– Blodgett troughs (LBT) is widely regarded as the benchmark technique for fabricating highly ordered colloidal monolayers for plasmonic sensors, offering excellent control over particle packing density and long-range order^28,29^. Nevertheless, LBT requires specialized instrumentation, precise control of interfacial pressure, and careful optimization of process parameters, which increases fabrication complexity and limits accessibility for routine sensor manufacturing^21^.

To address these limitations, we employ nanosphere settling lithography (NSSL) as a plasmonic sensor fabrication strategy that enables controlled interfacial assembly and transfer of colloidal monolayers using bottom-driven drainage under quasi-static conditions^30–32^. NSSL preserves the advantages of interfacial self-assembly while significantly reducing process complexity, equipment requirements, and sensitivity to experimental parameters. We demonstrate that NSSL produces large-area plasmonic sensor arrays with structural order, packing density, and optical uniformity comparable to LBT, while offering improved scalability. Using hyperspectral LSPR imaging, we demonstrate spatially uniform LSPR features and refractive-index sensitivity across 60 mm × 24 mm substrates compatible with multiwell biosensing formats. We further validate sensor performance through PANORAMA-based digital single-nanoparticle counting and single-vesicle analysis of antibody captured, tetraspanin-positive sEVs derived from cancer cell lines. Owing to its label-free detection capability, PANORAMA provides an assessment of captured particles with size-resolved information, while complementary fluorescence imaging enables quantitative analysis of the miR-21-positive fraction among captured sEV-like particles in patient plasma samples. Collectively, these results establish NSSL as a scalable, economical, and sensor-optimized fabrication approach for next-generation plasmonic biosensors and high-throughput digital single-sEV analysis.

## Experimental Section

### Materials

All reagents were purchased commercially and used without further purification. No. 1 glass coverslips were obtained from VWR. Aqueous PSB suspensions (0.46 µm, 2.5% w/v), neutravidin, buffered HF improved, and SDS were purchased from Sigma-Aldrich. The 460 nm PSBs were used exclusively as fabrication masks during the nanofabrication process. Additional PSB sizes (25, 50, and 100 nm), [MT(PEG)₄], and BSA were obtained from ThermoFisher Scientific. Biotin-PEG-thiol was purchased from Nanocs, and CD9, CD63, and CD81 antibodies from BioLegend. Ethanol (200 proof) was from Decon Laboratories. The 25 nm, 50 nm and 100 nm PSBs were used as standard particles for validating the PANORAMA imaging and detection capability. Titanium and gold sputtering targets (>99.99%) were from ACI Alloys. High-purity oxygen and argon gases (99.99%) were used for plasma etching and ion milling. DI water was obtained from a Milli-Q system. sEVs were extracted from 5 mL of H460 lung cancer cell culture medium (2–4 × 10⁶ cells/mL, ATCC).

## Methods

### Fabrication of AGNIS

An 80 nm thick gold film with 2 nm chromium adhesion layer was deposited on cleaned glass coverslips by sputtering. Thoroughly cleaned polystyrene beads (PSBs) of 460 nm diameter were assembled on the gold-coated substrates using three approaches: the petri dish method, Langmuir– Blodgett trough (LBT), and nanosphere settling lithography (NSSL) (See full details in Supplementary Note 1). Following deposition, oxygen plasma etching was employed to reduce the PSB diameter from 460 nm to approximately 360 nm. Argon ion milling was subsequently performed to selectively remove the exposed gold regions unmasked by the PSBs, yielding an arrayed gold nanodisk (AGN) structure beneath the beads. The PSBs were then removed by clamping the glass coverslip with a silicon wafer and subjecting the assembly to sonication in water. Finally, the nanodisk-patterned substrate was immersed in buffered hydrofluoric acid to undercut the glass beneath the disks, resulting in the formation of arrayed gold nanodisks on invisible substrates (AGNIS). This term “invisible” refers to the undercut dielectric support beneath the gold nanodisks, which significantly reduces substrate induced plasmonic coupling and minimizes the optical influence of the supporting substrate on the localized surface plasmon resonance response. The full details of the fabrication process are shown in Supplementary Note 1.

### Optical setup

#### LSPR mapping

Hyperspectral imaging of AGNIS was performed using an inverted Olympus microscope (IX71). A 100W white halogen lamp, focused with a 0.55 NA condenser, illuminated the AGNIS on the sample holder. The transmitted light was collected using a 40×/0.75 numerical aperture (NA) objective lens (UPlanFLN, Olympus) and directed to a spectrometer (Princeton Instruments, Acton 2300) with a thermoelectrically cooled (-70 °C) CCD camera (Princeton Instruments, PIXIS 400) through a telescope (L1 and L2). The optical setup achieved a spatial resolution of 2.5 µm, when measuring the AGNIS extinction spectrum. A schematic of the setup is provided in Supplementary Figure. S1a (Supplementary Note 2).

#### PANORAMA

The PANORAMA optical setup was built upon a commercial inverted microscope (Olympus IX83). Illumination was achieved using a tungsten-halogen lamp (U-LH100L3, Olympus), whose light was spectrally filtered using a bandpass filter (FSH660-10, Thorlabs) to narrow the wavelength range. This filtered light was then directed through a condenser (IX2-LWUCD, Olympus), enabling uniform illumination of the AGNIS substrate. Transmitted light was captured using an infinity-corrected air objective (UPlanSApo 40X/0.95, Olympus) and subsequently imaged with a scientific CMOS (sCMOS) camera (C14440-20UP ORCA-Fusion, Hamamatsu). A schematic representation of the optical setup is provided in Supplementary Figure S1b (Supplementary Note 2).

### Working Principle of PANORAMA

PANORAMA functions as a narrow-band bright-field imaging technique in which nanoparticle detection is achieved through transmission contrast generated by LSPR perturbations. Imaging is performed at a wavelength positioned near the half-maximum on left shoulder of the AGNIS LSPR extinction spectrum, where transmission is highly sensitive to resonance shifts. As a nanoparticle approaches the surface, the local dielectric environment is modified, resulting in a redshift of the LSPR. This resonance shift increases optical transmission at the selected wavelength, creating a localized enhancement in transmitted intensity beneath the particle.

### Image Processing for PANORAMA and Fluorescence

For each measurement location, multiple image frames (N= 50) were first averaged to suppress random noise. Images containing particles were then interpolated (1400×1400 pixels to 4000×4000 pixels) and aligned to corresponding background images acquired in the absence of particles. To correct for illumination non-uniformity, each particle image was normalized by dividing it by its background image. Particle detection was based on an intensity threshold defined as three times the background noise level plus a fixed offset (3σ + 0.5%). Assuming Gaussian noise behavior, signals exceeding this threshold were attributed to the presence of particles. Individual particles were identified by locating local intensity maxima, ensuring that each particle was counted only once. The identified particle centroids were then rendered as fixed-width Gaussian spots to generate a super-resolution style localization map, enabling clear visualization of diffraction-limited particle distributions with enhanced contrast while preserving their spatial positions.

Fluorescence images were processed in a similar way. Each fluorescence image was aligned to a background image and then corrected by subtracting the background signal. The corrected images were described by an average intensity and its variation. A detection threshold was set as three times the noise level above the average intensity. Based on Gaussian statistics, fluorescence signals above this threshold were assigned to particles containing miRNA. Similarly, like PANORAMA, the identified particle centroids were then rendered as fixed-width Gaussian spots. A detailed description of the image-processing workflow is provided in Supplementary Note 3.

### Surface functionalization of AGNIS

The surface modification of the AGNIS substrate was conducted in three sequential steps. Initially, the AGNIS substrate was incubated in a water solution comprising a 1:3 mixture of long (MW, 1 kDa) biotin-PEG-thiol and short (MW, 0.2 kDa) methyl-PEG-thiol polymers at 4°C for 16 hours to facilitate the binding of the thiol groups with the gold surfaces. In the second step, the modified substrate was incubated by a water solution of 3.3 µM neutravidin at 4°C for 2 hours. Finally, the substrate was incubated in a water solution containing 0.5 mg/mL of biotinylated CD9, CD63, CD81 antibodies, and 2.5% BSA at 4°C for 2 hours to allow for antibody modification. The resulting AGNIS chip was utilized for sEV detection experiments.

### Plasma Sample Collection and Processing

Use of de-identified samples for sEV analysis was conducted under IRB protocol 2021-0368 (ID: 2021-0368_CR001; IRB 2 IRB00002203). Peripheral blood samples were collected via venipuncture prior to the initiation of any therapeutic intervention, including radiation therapy, surgery, or chemotherapy. Following collection, blood samples were processed in the laboratory to isolate plasma. The extracted plasma was aliquoted and stored at −80 °C until further analysis. All samples used in this study were de-identified prior to downstream processing. All procedures were performed in compliance with institutional guidelines and applicable ethical standards.

### sEV isolation from cancer cell line cultures

H460 cells were initially cultured in RPMI1640 supplemented with 10% FBS, then switched to exosome-free medium for 48 h. The conditioned medium was cleared of debris by centrifugation at 2,000 × g for 10 min, followed by removal of large vesicles and apoptotic bodies at 10,000 × g for 30 min. sEVs were concentrated by centrifugation at 3000 × *g* for 30 min using Amicon® Ultra centrifugal filters (Sigma UFC9100) and further purified with Total Exosome Isolation Reagent (Thermo Fisher 4478359) per manufacturer instructions. Purified sEVs were resuspended in cold PBS.

### Molecular beacon probe design and delivery into sEV

A Cy3-labeled molecular beacon probe complementary to miR-21 (5′Cy3/GCGCGTCAACATCAGTCTGATAAGCTACGCGC/3′BHQ; Integrated DNA Technologies) was used to detect intravesicular microRNA (*miR-21*). The probe possesses an initial hairpin structure that positions the Cy3 fluorophore in close proximity to the black hole quencher (BHQ), thereby suppressing fluorescence in the absence of target binding. A 250 nM working solution of the *miR-21* molecular beacon was incubated with 20 µL of plasma sample under light-restricted conditions to minimize photobleaching. The mixture was incubated at 40 °C for 2 h to facilitate molecular beacon permeation into sEVs and subsequent hybridization with target *miR-21* sequences. Upon hybridization, the molecular beacon undergoes a conformational transition to an extended structure, separating the Cy3 fluorophore from the BHQ quencher. This spatial separation restores fluorescence emission, thereby confirming target binding within the vesicles.

## Results and discussion

### Structural and Optical Characterization of AGNIS

High-resolution electron microscopy was employed to evaluate PSB ordering and pattern transfer fidelity in AGNIS substrates. Spatial non-uniformities arising during PSB monolayer assembly were observed to be directly transferred into the final nanodisk array following pattern transfer. Figure 1a shows a 60 mm × 24 mm AGNIS substrate fabricated using the NSSL method, sufficient for 60 wells in a standard 384-well plate format, with 4 substrates accommodating up to 240 sensing wells. Quantitative SEM-based analysis was performed to determine nanodisk diameter distributions and interdisk spacing. A representative SEM image of the AGNIS substrate is shown in Figure 1a, with the corresponding nanodisk diameter distribution presented in Figure 1b, yielding an average diameter of 359 ± 7.5 nm. The center-to-center spacing between adjacent nanodisks was measured to be 433 ± 20 nm (Figure 1c). The relatively narrow distributions in both nanodisk diameter and interdisk spacing observed indicate the formation of densely packed and spatially uniform monolayers, confirming that the NSSL approach yields uniformly packed hexagonal AGNIS arrays.

**Figure 1:**
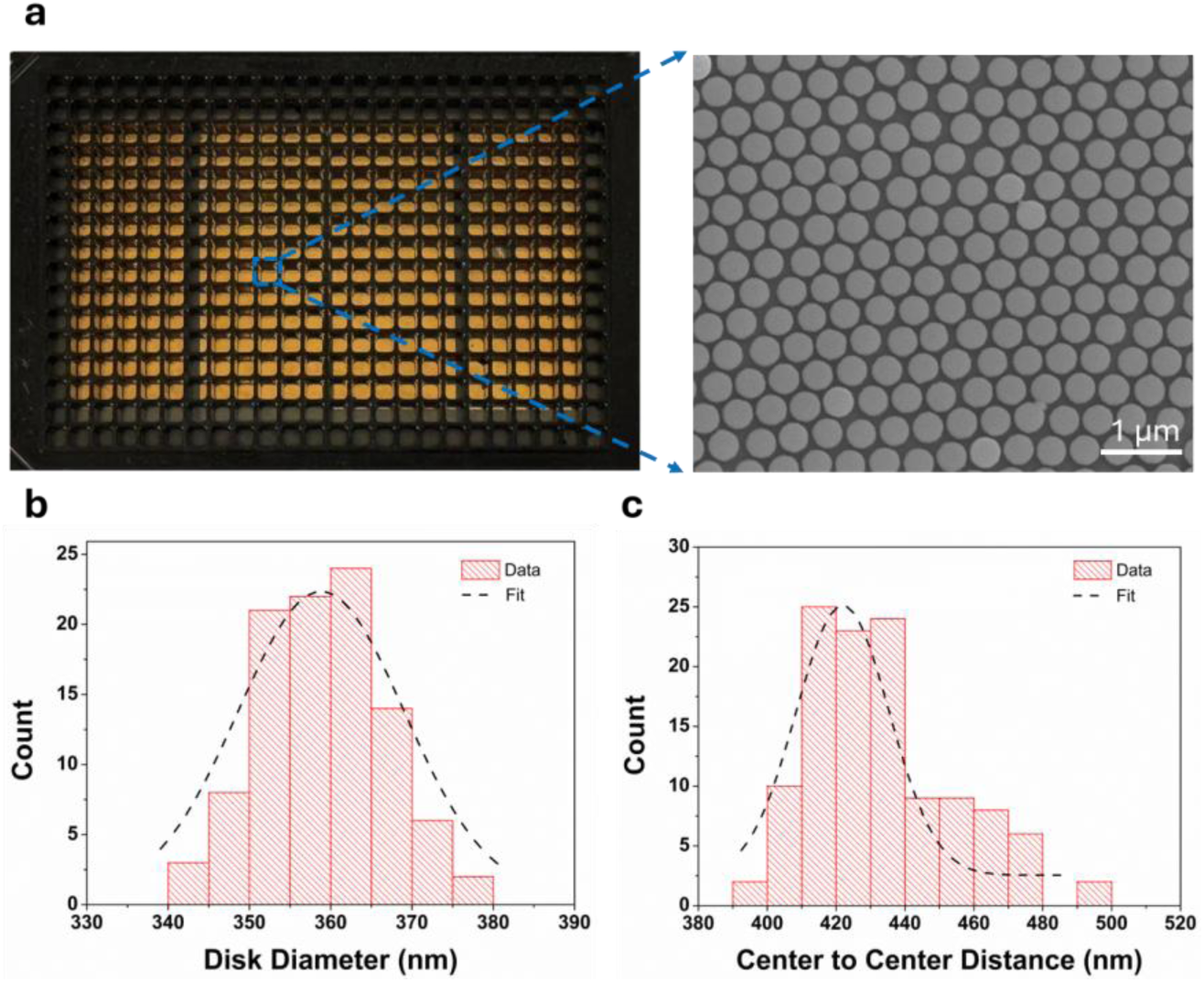
**a.** Four 60 mm × 24 mm AGNIS fabricated via NSSL method accommodating 240 wells in the 384 well plate format and SEM image of the fabricated AGNIS. Distribution of the **b.** diameter of nanodisks and **c.** center to center distance in AGNIS. The diameter and center to center distance were calculated from 5 different images (each 10 µm x 10 µm).

The reproducibility and reversibility of the refractive-index-dependent LSPR sensing response was assessed by measuring peak shifts across multiple wells (N = 6) during five successive refractive index modulation cycles using air and water (Figure 2a). A mean LSPR peak shift of 120.5 nm was observed between air and water environments, yielding a sensitivity of 365 ± 2 nm/RIU. This value represents the average refractive-index sensitivity obtained from repeated air-to-water switching measurements performed at the same sensing location. Hexagonal packing uniformity directly influences the spatial uniformity of localized surface plasmon resonance (LSPR) through variations in nanodisk dimensions and interparticle coupling. Structural disorder can therefore introduce spatial variability in refractive index sensitivity. To quantify optical uniformity, hyperspectral imaging was employed to map the LSPR response over a 100 µm × 200 µm region with a spatial resolution of approximately 2.5 µm. The LSPR sensitivity map (mean ± σ) (Figure 2b) yielded average peak positions of 640 ± 4 nm and 758 ± 6 nm, respectively, indicating a pronounced refractive-index-induced spectral shift accompanied by a narrow spatial distribution. Corresponding histograms derived from hyperspectral imaging data (Figure 2c) yielded an average refractive index sensitivity of 356 ± 21 nm/RIU, where the mean and standard deviation represent the spatial distribution of sensitivity across the mapped sensor region. Additional measurements conducted at five distinct locations on a single coverslip (Figure 2d) demonstrated consistent sensitivity across inch-scale distances, confirming excellent optical reproducibility and large-area uniformity.

**Figure 2:**
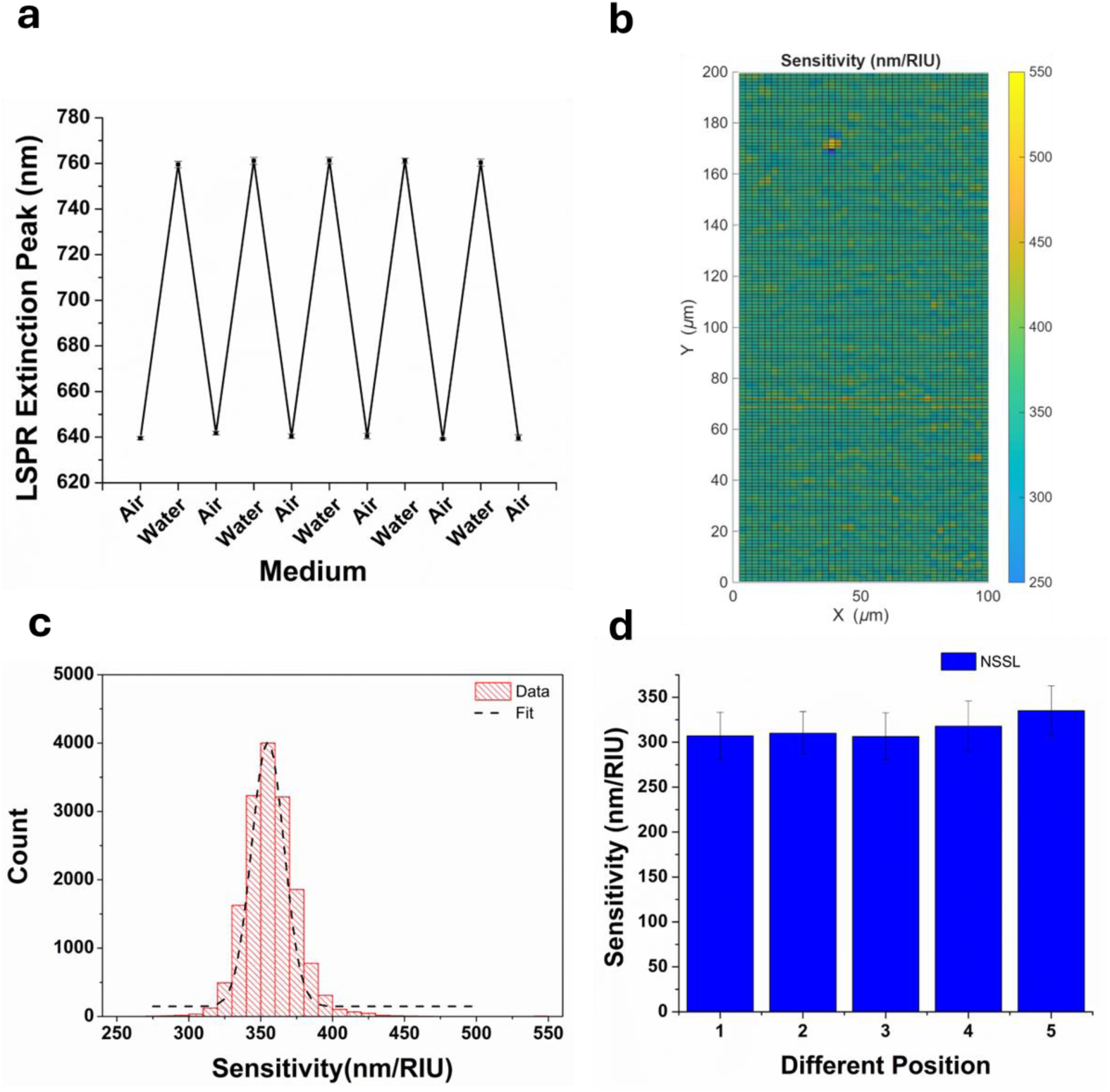
**a.** LSPR peak shift for alternating air and water. The error bars represent the standard deviation of the plasmonic response from 6 wells each measured 8 times per environment**. b**. Spatial mapping of the refractive-index sensitivity of AGNIS. **c**. Distribution of the refractive index sensitivities obtained from experimentally measured extinction spectra (mean±std) of AGNIS over an area of 200×100 µm^2^. **d.** Comparison of the mean refractive index sensitivity of several different regions (N= 5) on a single AGNIS. The error bars represent the standard deviation of 40 measurements with a spatial resolution of approximately 2.5 µm.

### PANORAMA digital single-nanoparticle analysis

PANORAMA leverages the optical properties of AGNIS to achieve high sensitivity detection of nanoscale particles potentially down to below 10 nm^12^ (see Methods for details of the detection mechanism and image-processing workflow). PANORAMA images of detecting PSBs of 25 nm, 50 nm, and 100 nm in diameter are shown in Figure 3a–c, taken at 60 minutes after dispensing 20 microliters of PSB solutions at a concentration of ∼10^6^ particles/mL. Most PSBs appeared as patches of 3×3 pixels, with the highest contrast in the center pixel, surrounded by pixels of lower contrast. PANORAMA mean contrasts values for the three different sized PSB targets were measured to be 10.6±0.4%, 14.9±0.5%, and 17.7±1% for 25 nm, 50 nm, and 100 nm PSBs, respectively (Figure 3d) across 3 repetitions. Among the 3 repeats, the PSB counts (mean ± σ) were 188±5, 174±15, and 143±14 for 25 nm, 50 nm, and 100 nm PSBs, respectively (Figure 3e).

**Figure 3:**
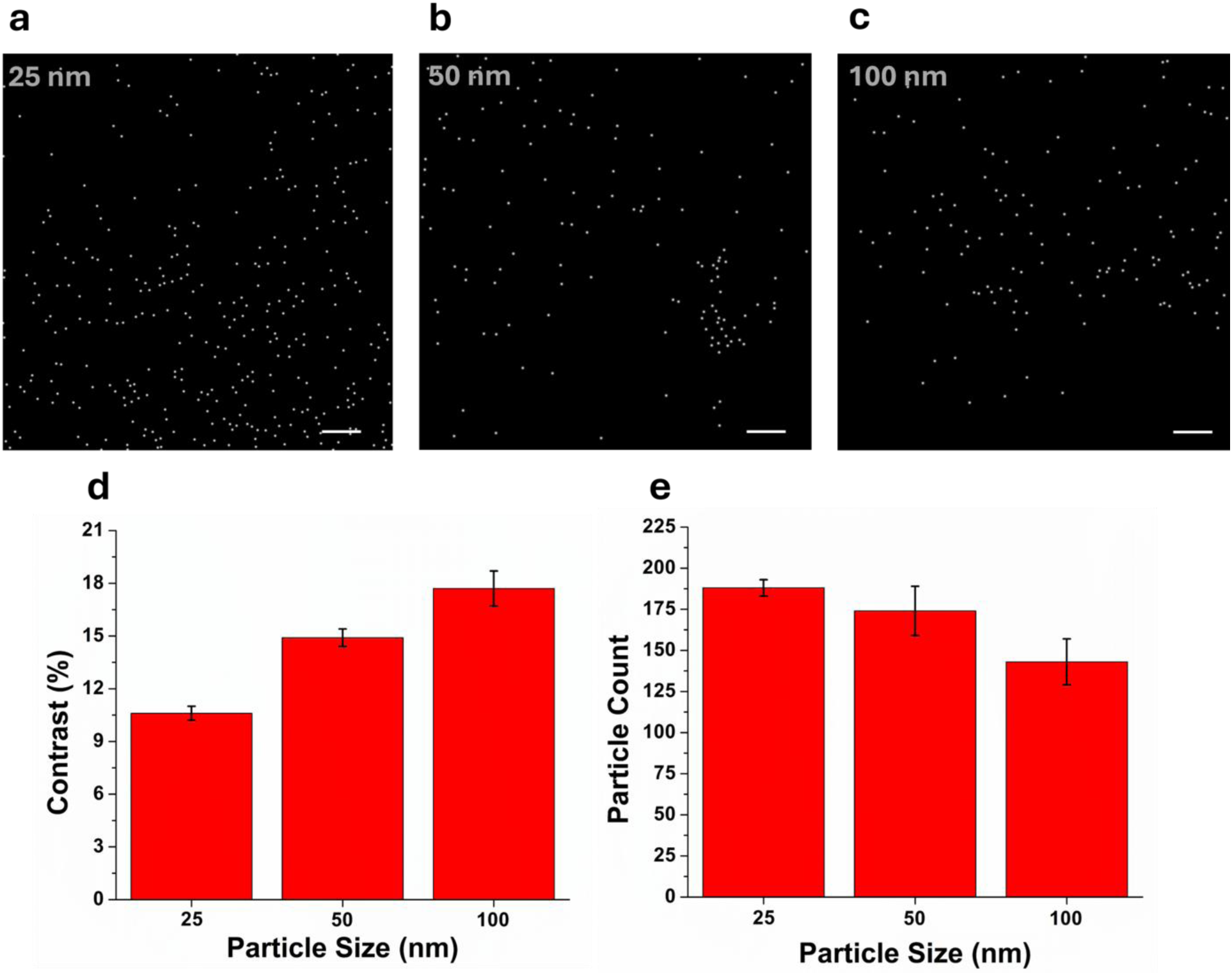
PANORAMA images of detected PSBs **a.** 25 nm **b.** 50 nm **c.** 100 nm with respective **d.** contrast and **e.** counts with three chips (N= 3). Scale bar: 10 μm.

SEM analysis further confirmed a one-to-one nanoparticle-to-nanodisk capture configuration, with more than 90% of nanodisks remaining unoccupied. SEM-based particle occupancy analysis showed that the distribution of particles across individual AGNIS nanodisks was consistent with a Poisson distribution (Supplementary Note 4). This is a salient feature showcasing single particle detection where overwhelmingly abundant sensing sites are available. Additionally, the large-area substrate configuration (60 mm × 24 mm; Figure 1a) enabled up to 240 (60 × 4) experiments to be performed in parallel, demonstrating the scalability and suitability of this fabrication approach for high-throughput PANORAMA measurements. To demonstrate real-time single particle monitoring capability, a video was recorded following injection of the particle solution, during which individual 100 nm PSBs appeared on the AGNIS surface over time (Supplementary Video S1). Direct sensing measurements performed under identical assay conditions showed that NSSL-fabricated AGNIS substrates produced the highest mean particle count compared with the petri-dish and LBT methods (Supplementary Note 5), supporting the suitability of NSSL for scalable and robust biosensor fabrication.

### PANORAMA digital single-small extracellular vesicle (sEV) from H460 cancer cell line cultures

Biosensing experiments were conducted to analyze purified sEVs secreted by the H460 lung cancer cell line. The sEV extraction protocols are provided in Methods. The same protocols have been proven to provide high purity sEVs without protein aggregates using PKH67 membrane bound fluorescent markers^19^. To enable effective and specific surface capture, AGNIS substrates were functionalized with antibodies targeting the sEV surface markers CD9, CD63, and CD81 according to protocols described in Methods. The stepwise surface functionalization process was monitored using hyperspectral imaging by tracking changes in the (LSPR) peak. AGNIS substrates were first functionalized with a mixed self-assembled monolayer (SAM) consisting of biotinylated thiol-PEG and methyl-terminated (inactive) thiol-PEG for 16 h, followed by PBS washing. This modification induced a redshift in the LSPR peak from 757 nm (PBS baseline) to 768.7 nm, confirming successful PEG layer formation (Figure 4a). Subsequent incubation with 3.3 µM neutravidin for 2 h, followed by washing, produced an additional peak shift to 775.8 nm, consistent with protein adsorption via specific biotin–neutravidin interactions. Next, 0.5 mg mL⁻¹ biotinylated anti-CD9, anti-CD63, and anti-CD81 antibodies were immobilized in the presence of 2.5% BSA. After washing, a further redshift to 779.8 nm was observed, confirming successful antibody attachment to the surface. Finally, exposure to sEVs at a concentration of 1 × 10⁷ sEVs mL^-1^ resulted in a final peak position of 786.7 nm after washing, indicating successful vesicle capture through antibody–antigen binding (Figure 4a). The cumulative LSPR peak shifts at each functionalization step are summarized in Figure 4b across multiple wells (N = 8), demonstrating consistent and reproducible surface modification and target binding.

**Figure 4:**
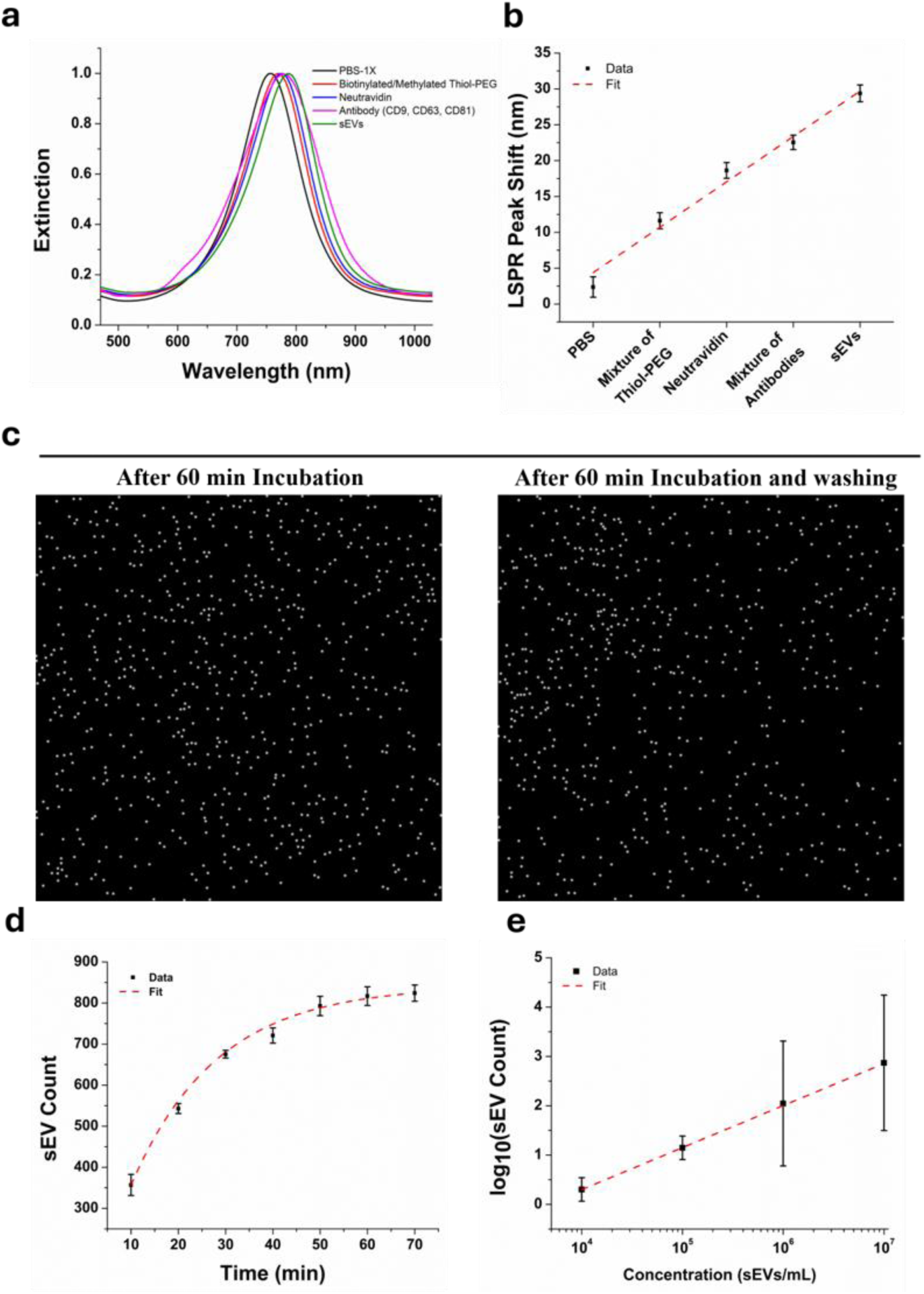
**a.** Monitoring of surface functionalization steps using hyperspectral imaging. **b.** Successive LSPR peak shift after each step. **c.** PANORAMA images of detected sEVs isolated from H460 cancer cell line cultures. **d.** Time-dependent sEV count obtained from three independent experiments. **e.** Captured sEV counts vs. concentrations. Scale bar: 10 μm.

For PANORAMA experiments, a 20 µL sample of purified sEVs (10⁷ sEVs mL^-1^) was applied to the functionalized AGNIS substrates and monitored for 60 min, followed by washing. Representative PANORAMA images showing detected sEVs before and after washing 5 times after 60 min are presented in Figure 4c, where ∼11% of particles were removed. The retained sEVs are observed in SEM images (Supplementary Figure S5c). A video recorded prior to sample injection showed no particles on the AGNIS surface. Upon injection of the sEV solution, particles started to appear, confirming real-time vesicle detection (Supplementary Video S2). The washing protocol was optimized to remove unbound or weakly adhered vesicles while preserving specifically bound sEVs. AGNIS chips were washed with PBS-1X for 0, 3, 5, 10, and 15 cycles. Successive removal of particles persisted until the five wash cycles, beyond which the detected sEV count remained mostly constant (Supplementary Note 6), indicating stable and specific capture.

Figure 4d shows the time-dependent sEV counts within 60 minutes, demonstrating monotonic accumulation over time. The temporal evolution of particle counts was modeled using a first-order exponential function to describe the progressive occupation of available surface binding sites over incubation time (R^2^ ≈ 0.9949). The extracted characteristic time constant was 19.24 ± 2.43 min with a corresponding effective binding rate constant 0.0520 ± 0.0035 min^-1^. A weighted nonlinear least-squares fit using the inverse variance of the measured standard deviations (1/σ^2^) yielded an an asymptotic count at equilibrium of 848.39 ± 23.22, suggesting ∼95% of the asymptotic count was achieved within 57.63 min. NTA measurements confirmed the particle size distribution and concentration of the samples (Supplementary Note 7), supporting characterization of the tetraspanin-captured sEV particle population analyzed in this study.

To further confirm that individual nanodisks predominantly capture single sEVs, SEM imaging was performed following vesicle fixation using a standard protocol^33^, revealing a one-to-one sEV-to-nanodisk capture configuration (Supplementary Figure S5c). Since only about 850 sEVs are detected within the 200 x 200 μm^2^ area which houses ∼2.18×10^5^ nanodisks, the sEV per nanodisk is 0.4% statistically, highlighting that assay performance is governed primarily by effective capture and detection efficiency rather than total geometric surface area. Additional experiments were conducted by varying the concentration of H460-secreted sEVs across four orders of magnitude, and the results are shown in Figure 4e. The achievable concentration limit of detection (LOD) was 1 × 10⁴ sEVs mL^-1^, corresponding to approximately 200 particles per well or 17 aM with a broad dynamic range. The LOD was determined experimentally by performing sensing experiments with serial 10-fold dilutions of the sEV suspension and defined as the lowest concentration at which sEVs could be reliably detected above the background signal. A quantitative comparison of the LOD and response time of representative sEV detection technologies is provided in Supplementary Table S1(Supplementary Note 8). To further assess detection specificity at low concentrations, control experiments were performed using biotinylated IgG in place of anti-CD9/CD63/CD81 antibodies for surface capturing. No detectable binding was observed at the lowest concentration, while only minimal nonspecific binding (0.9%) was detected at the highest concentration (Supplementary Figure S5d), confirming the specificity of antibody-mediated sEV capture on the AGNIS substrate.

### PANORAMA and fluorescent imaging of sEV *directly* from human blood plasma samples without isolation or purification

PANORAMA has been demonstrated to provide label-free digital single-sEV detection after selective anti-CD9/CD63/CD81 affinity capture. Next, we would like to further interrogate the molecular contents in individual sEVs by fluorescent probes. In the literature, micronRNA-21 (*miR-21*) has been identified among sEVs associated with several types of cancer. Joint label-free and labeled analysis at single-sEV level over the same antibody captured sEV-like particle population is important for identifying the miR-21-positive fraction among captured particles. We next evaluated the dual-modality capability of our technique to detect and quantify sEVs directly from the plasma of a liver cancer patient without prior isolation and purification. Plasma samples (20 µL) were dispensed into the antibody-functionalized AGNIS well, incubated for 60 min, and subsequently washed to remove nonspecifically bound entities. Following the washing step, co-registered PANORAMA and fluorescence images were acquired. For fluorescence imaging, *miR-21* was targeted using Cy3-labeled molecular beacon probes which only emit fluorescence light after the probe hybridizes with *miR-21* sequence. The specific molecular beacon probe design and a protocol of delivering them into sEVs were developed and validated in our previous paper, and the details are provided in Methods. Specificity controls using BSA-blocked surfaces and sEV-depleted plasma showed substantially lower PANORAMA-positive particle counts than anti-CD9/CD63/CD81-functionalized surfaces (Supplementary Note 9). These findings indicate that the assay preferentially detects a tetraspanin-captured, sEV-enriched nanoscale particle population rather than nonspecific adsorption alone. To validate the specificity of the fluorescence assay, we performed additional control experiments using sEV-depleted plasma, plasma without miR-21, miR-21 molecular beacon alone, plasma with miR-21 probes, and molecular beacon delivery with and without mild heating. These control experiments support our interpretation that the detected fluorescence originated from intravesicular miR-21 in captured sEVs rather than nonspecific adsorption or free nucleic acids (Supplementary Note 9 and Supplementary Figure S7–S9).

Figure 5a-b shows PANORAMA, fluorescence, and their merged images of sEVs detected at 60 min post wash, which allows us to distinguish three categories of sEV populations: PANORAMA positive, Fluorescence positive (P^+^F^+^, 74), PANORAMA positive, Fluorescence negative (P^+^F^-^, 585), and PANORAMA negative, Fluorescence positive (P^-^F^+^, 17). A detailed description of the image-processing workflow is provided in Supplementary Note 3. The experiment and corresponding analysis were repeated three times, demonstrating good reproducibility with a coefficient of variation (CV) of ∼3.46% for the total sEV counts (mean ± σ) of 695 ± 24 with a split of 65 ± 13 for the P^+^F^+^ population and 611 ± 23 for the P^+^F^-^ population (Figure 5c). The *miR-21* expression level at the single-sEV level is thus 9.7±1.9% (=P^+^F^+^/(P^+^F^+^+P^+^F^-^)), indicating a significant portion of sEVs carry *miR-21* as observed in the literature^19,34–36^. Within the antibody captured PANORAMA-positive population, the P⁺F⁺ fraction was interpreted as the proportion of captured sEV-like particles containing detectable miR-21. In contrast, the P⁺F⁻ fraction should not be considered definitively cargo-negative, as it may include sEVs lacking the fluorescent target, vesicles with target abundance below the fluorescence detection limit, disrupted vesicles, or residual non-vesicular plasma particles. Since the binding status of the P⁻F⁺ population cannot be confirmed and this population accounted for only 17 particles, approximately 2% of the total detected particles, we excluded these events from the quantitative analysis. These interpretations highlight the unique capability of our dual-modality technique to achieve both affinity captured sEV detection and molecular profiling directly from complex clinical plasma samples.

**Figure 5:**
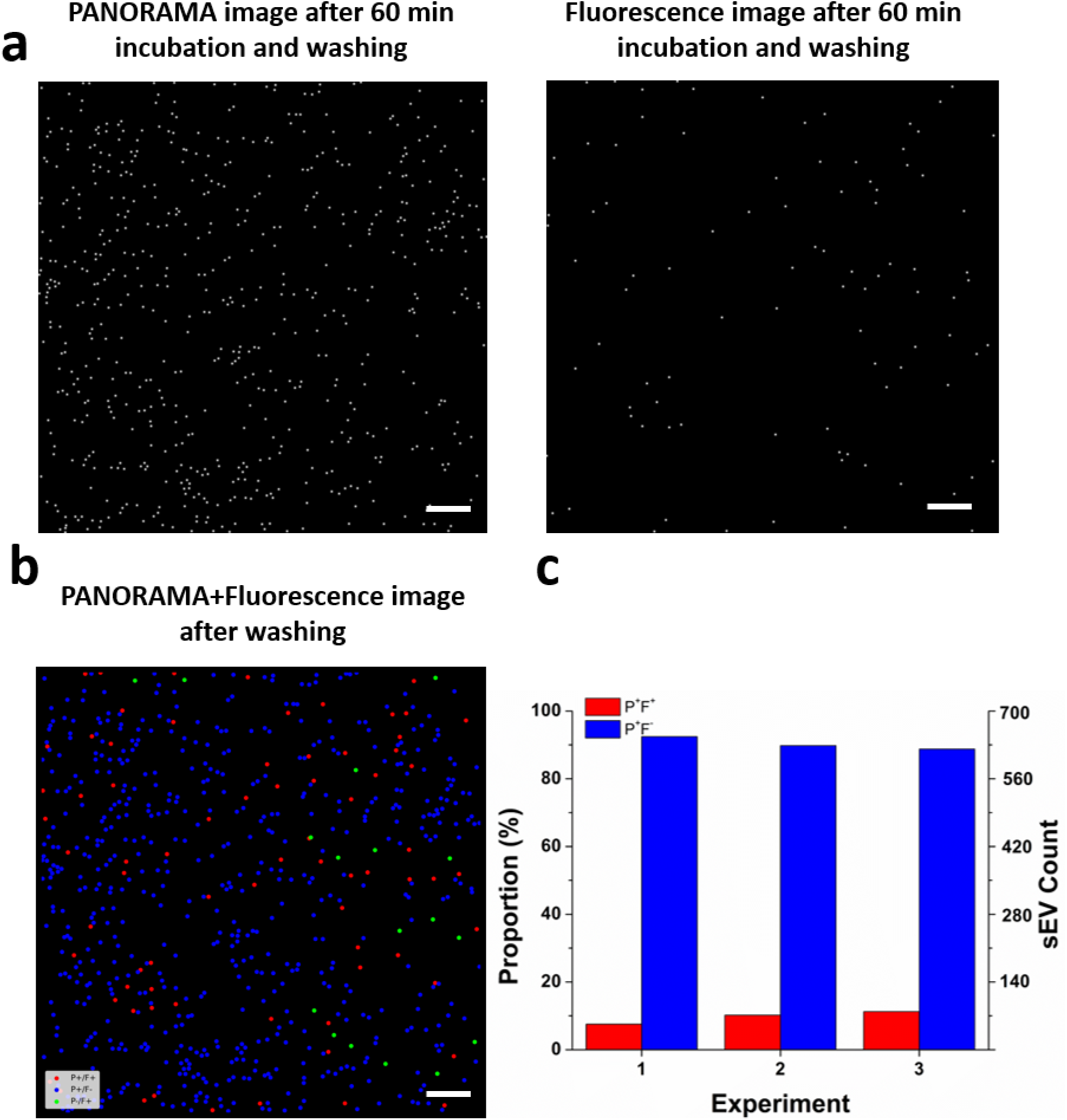
**a.** PANORAMA and fluorescence images of sEVs from liver cancer plasma samples. **b.** Merged PANORAMA and fluorescence image of sEVs. **c.** Count and proportion split for P^+^F^+^ and P^+^F^-^ sEVs with three repetitions. Scale bar: 10 μm.

## Conclusion

In conclusion, we have demonstrated isolation and purification-free digital antibody-captured single-sEV analysis on scalable AGNIS arrays fabricated by nanosphere settling lithography (NSSL). The densely packed hexagonal AGNIS arrays exhibit tight distribution of nanodisk size and spacing, resulting in uniform LSPR responses and an average refractive index sensitivity of 356 ± 21 nm/RIU across the substrate. The uniform sensitivity enables highly reproducible label-free digital sEV detection from H460 cell cultures with a LOD of 1×10^4^ sEVs mL^-1^ or 17 aM. Combining PANORAMA and fluorescence imaging enables quantitative analysis of the miR-21-positive fraction among captured sEV-like particles from liver cancer patient plasma, demonstrating the potential of this dual-modality approach for molecular profiling of affinity captured nanoscale vesicle populations. AGNIS thus fabricated exhibit high refractive-index sensitivity with low spatial variability across 60 mm × 24 mm substrates, sufficient for 60-well in standard 384-well plate format. A high-throughput platform of 240 experiments in parallel is envisioned by assembling 4 AGNIS with a 384 well plate.

## Author Contributions

W.-C.S. conceived the idea, obtained funding to support this research, and directed the study. S.M. and M.S.M fabricated the AGNIS. M.S.M, S.M., A.K., and A.H. acquired and analyzed all the experimental data. M.S.M., S.M., A.K., and W.-C.S. interpreted the data and explained the results and wrote the manuscript.

## Acknowledgements

Cell line sEVs were provided by Dr. Steven H. Lin (MD Anderson Cancer Center), and deidentified liver cancer plasma samples were provided by Dr. Manal M Hassan (MD Anderson Cancer Center) and Dr. Prasun Jalal (Baylor College of Medicine).

## Data availability statement

All details of the data analysis are provided in the Supplementary Information. A web link to the datasets will be made available upon request.

## Supplementary Information

Supporting Information Available: The following files are available free of charge.

File name: Supplementary Information, Supplementary Video S1, and Supplementary Video S2.

Supplementary Information includes additional experimental procedures, AGNIS fabrication and characterization including details of Poisson analysis of particle occupancy on AGNIS, image processing workflow and a series of control experiment data supporting the conclusions of this work.

