## Supplementary material for "Isolation and Purification-Free Digital Single-Small Extracellular Vesicle Biosensing with Scalable Plasmonic Arrays": Suppl. Info

**Supplementary Note 1:**

**Preparation of Polystyrene Beads and Substrates**

Polystyrene beads (PSB) were first cleaned to remove surfactants and increase hydrophilicity. The beads were rinsed in a water–ethanol (1:1) mixture, followed by 6–7 centrifugation cycles and drying. The cleaned beads were then dispersed in a water–ethanol (1:2) solution by adding 100 µL of solvent per 1 mg of PSB.

Glass coverslips were cleaned sequentially in acetone and isopropanol baths, rinsed with copious DI water, and dried with nitrogen. A 2 nm Cr adhesion layer and 80 nm Au film were deposited by sputtering.

**Petri Dish Monolayer Assembly**

Approximately 250 µL of 460 nm PSB solution was dispensed onto the water surface using a syringe pump. This formed a white, turbid film at the air-water interface, indicating that the water surface was fully covered with a PSB monolayer. A drop of surfactant (sodium dodecyl sulfate) solution was added to a corner of the petri dish, which helped bring the PSB beads closer together. The substrate was then manually lifted horizontally using tweezers. Finally, the transferred film was dried on a hot plate at 70°C.

**Langmuir–Blodgett Trough (LBT) Method**

A commercial Langmuir-Blodgett trough (KSV NIMA) was employed. A monolayer film was formed by gently dispersing the PSB solution at the air-water interface using a syringe pump. Initially, the PSB particles moved randomly at the interface, gradually forming smaller monolayer domains. As more PSB solutions were added, the particles came closer together, creating a white, turbid film that indicated the water surface was fully covered with a PSB monolayer. The film was then compressed using robotic barriers, and surface pressure was monitored so that it doesn’t exceed 30 mN/m, promoting the formation of a continuous, hexagonally packed monolayer structure. The substrate, which had been previously immersed in the water inside the Langmuir-Blodgett trough, was then vertically pulled out, transferring the monolayer onto its surface while maintaining a constant surface pressure. Finally, the substrate was dried on a hot plate at 70°C to complete the process. The surface pressure was automatically regulated throughout the coating process using KSV NIMA software-controlled barriers to ensure uniformity.

**Nanosphere Settling Lithography (NSSL)**

A substrate was fully submerged in deionized (DI) water, followed by a similar setup to the petri dish method. Approximately 250 µL of PSB solution was then carefully dispensed onto the water surface using a syringe pump, creating a white, turbid film at the air-water interface. This film indicated the formation of a PSB monolayer. To improve the packing density of the polystyrene beads, a drop of sodium dodecyl sulfate (SDS) surfactant was added to one corner of the petri dish. This addition facilitated better organization of the beads by reducing surface tension. Gradual water removal via a drain at the petri dish bottom was initiated after the PSB monolayer had formed on the water surface. To regulate the drainage process, the hole was initially sealed with a transparent adhesive tape and then precisely punctured using a sharp pin, allowing controlled water removal. As the water level slowly lowered, the floating PSB monolayer gently settled onto the submerged substrate. The slow removal of water ensured a uniform deposition of the monolayer, minimizing defects during the transfer. Once the water was completely drained, leaving the PSB monolayer fully transferred onto the substrate, the coverslips were dried on a hot plate at 70°C to complete the NSSL process.

**PSB Monolayer to AGNIS Fabrication**

After the deposition of the 460 nm PSB monolayer onto the gold-coated glass coverslip substrates, oxygen plasma etching was employed to reduce the diameter to 360 nm. The etching parameters were set as follows: oxygen flow rate of 50 sccm, chamber temperature of 21°C, chamber pressure of 30 mTorr, and RF power of 50W. Argon milling was then performed to selectively remove the underlying gold unmasked by the PSBs, resulting in an arrayed gold nanodisk (AGN) under PSBs. The argon milling parameters included a normal angle of incidence, a sample holder distance of 30 cm, and an etch rate of 0.62 nm/s. The PSBs were then removed by clamping the glass coverslip substrate with a silicon wafer and the assembly was subjected to a 20-minute sonication process in water. After the removal of the PSBs, the gold nanodisks covered substrate was immersed in a buffered hydrofluoric acid solution for 85 sec to undercut the glass substrate beneath the disks, resulting in the final nanostructured plasmonic sensor, arrayed gold nanodisks on invisible substrates (AGNIS). This undercut structure effectively eliminates the substrate effect, resulting in AGN formed on an “invisible” substrate.

**Supplementary Note 2:**

The AGNIS chips fabricated were optically characterized by LSPR hyperspectral mapping and PANORAMA were performed using the optical setups described in Supplementary Figure S1a and 1b. Briefly, LSPR mapping was carried out on an inverted microscope coupled to a spectrometer for spatially resolved extinction spectra, and PANORAMA used a filtered halogen illumination path with a high-NA objective and sCMOS camera.

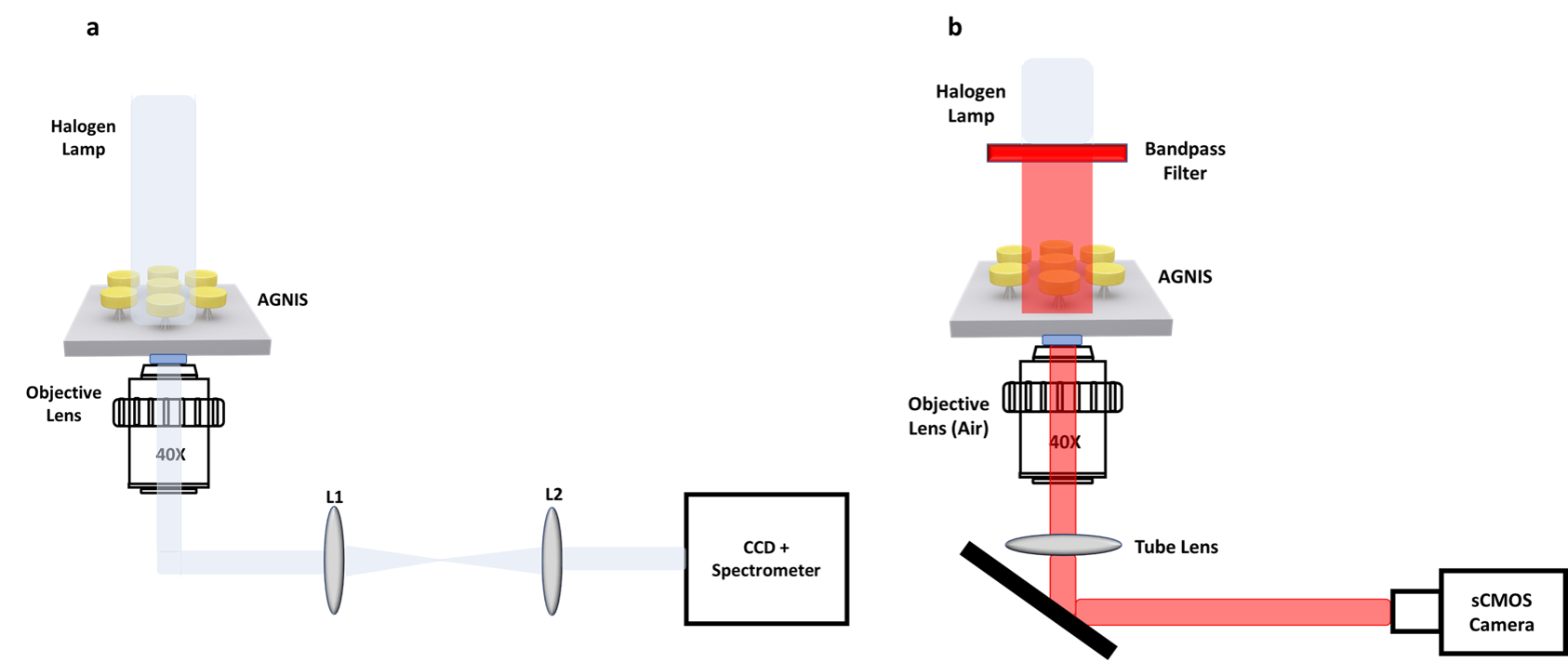

**Supplementary Figure S1.** Optical setup of **a.** LSPR mapping system **b.** PANORAMA.

**Supplementary Note 3:**

**PANORAMA Image Processing**

PANORAMA images were processed using ImageJ. For each measurement location, multiple sequential image frames (N= 50) were first averaged to reduce random camera noise and improve the signal-to-noise ratio. The averaged PANORAMA image (with bounded particles) was then interpolated (1400×1400 pixels to 4000×4000 pixels) and spatially registered to a corresponding background image acquired from the same sensing area prior to particle binding. Image registration minimized errors arising from mechanical drift or slight stage misalignment between acquisitions. To compensate for non-uniform illumination and variations in sensor response across the field of view, the registered particle image was normalized by dividing it pixel-by-pixel by its corresponding background image, producing a ratio image in which localized intensity changes associated with particle binding were enhanced while slowly varying background intensity was suppressed. An additional local mean intensity correction was subsequently applied to further compensate for residual illumination gradients that remained after normalization.

Particle detection was performed on the normalized ratio images using a threshold-based approach. The background noise level was estimated from particle-free regions of each image, and the detection threshold was defined as three times the standard deviation of the background noise (3σ) plus a fixed intensity offset of 0.5%. Assuming Gaussian statistics for background noise distribution, pixels exceeding this threshold were considered statistically significant particle signals.

Individual particles were identified using the ImageJ “Find Maxima” algorithm, which detects local intensity maxima corresponding to diffraction-limited particle images. The prominence parameter was selected to be greater than (Threshold −1), ensuring that only intensity peaks significantly above the surrounding background were identified while suppressing detections arising from random noise fluctuations. Large dust particles and imaging artifacts were excluded using an area-based filtering procedure.

The centroid coordinates of each detected particle were extracted and rendered as fixed-width Gaussian spots to generate a super-resolution–style localization map. This rendering provides a high-contrast visualization of the spatial distribution of diffraction-limited particles while preserving the experimentally determined particle positions without altering their measured locations.

**Fluorescence Image Processing**

Fluorescence images were processed using a custom MATLAB algorithm. For each measurement location, the fluorescence image was first spatially registered to a corresponding background image acquired under identical imaging conditions prior to sample introduction. The background image was subtracted from the fluorescence image to remove detector offsets, autofluorescence, and non-uniform background emission. The resulting background-corrected image was characterized by its mean background intensity and corresponding standard deviation. A detection threshold was then established as the mean background intensity plus three times the standard deviation (Mean + 3σ). Under the assumption of Gaussian-distributed background noise, fluorescence signals exceeding this threshold were classified as statistically significant and attributed to particles containing target miRNA.

Detected fluorescent particles were identified by locating local intensity maxima within the thresholded image, ensuring that each diffraction-limited fluorescence spot was counted only once. The centroid position of every detected particle was then calculated and represented as a fixed-width Gaussian spot to generate a super-resolution–style localization map. This approach enhances visualization of the spatial distribution of miRNA-positive particles while maintaining experimentally measured particle locations and enabling direct comparison with the corresponding PANORAMA localization maps.

**Classification of PANORAMA and Fluorescence Particle Populations**

Processed PANORAMA and fluorescence images were analyzed to classify particles into PANORAMA-positive/fluorescence-positive (P⁺F⁺), PANORAMA-positive/fluorescence-negative (P⁺F⁻), and PANORAMA-negative/fluorescence-positive (P⁻F⁺) populations. Particle detection was performed independently for each imaging modality. Images were converted into binary masks using an intensity threshold and connected-component analysis was applied to identify individual particle-associated regions. The centroid coordinates and projected areas of all detected particles were subsequently extracted for particle matching.

Spatial correspondence between PANORAMA and fluorescence particles was established using a nearest-neighbor search based on a KD-tree algorithm. Owing to differences in image formation, point spread function, and localization precision between the PANORAMA and fluorescence imaging modalities, the apparent particle patch size and centroid position can differ by up to approximately 10-12 pixels for the same physical particle. A distance-based nearest-neighbor matching criterion was therefore employed to identify corresponding particles while accounting for these modality-dependent localization offsets. Spatially matched particles were classified as PANORAMA-positive/fluorescence-positive (P⁺F⁺), with each fluorescence particle uniquely assigned to a single PANORAMA particle to prevent duplicate matching.

PANORAMA particles that did not have a corresponding fluorescence particle were classified as PANORAMA-positive/fluorescence-negative (P⁺F⁻), whereas fluorescence particles that remained unmatched after the nearest-neighbor matching procedure were classified as PANORAMA-negative/fluorescence-positive (P⁻F⁺). To minimize false-positive fluorescence detections arising from isolated noise pixels or small imaging artifacts, unmatched fluorescence objects with projected areas smaller than four pixels were excluded from further analysis.

The algorithm quantified the total numbers of PANORAMA-positive particles, fluorescence-positive particles, and the P⁺F⁺, P⁺F⁻, and P⁻F⁺ populations for each image pair. The classified particles were subsequently overlaid on the processed PANORAMA image using distinct color-coded markers to visualize particle co-localization between the PANORAMA and fluorescence imaging modalities.

**Supplementary Note 4:**

**Poisson Analysis of Particle Occupancy on AGNIS Nanodisks using SEM Images**

SEM imaging, performed at an accelerating voltage of 15 kV after sample drying and Au coating for conductivity, confirmed the binding of 100 nm PSBs to AGNIS (Supplementary Figure S2).

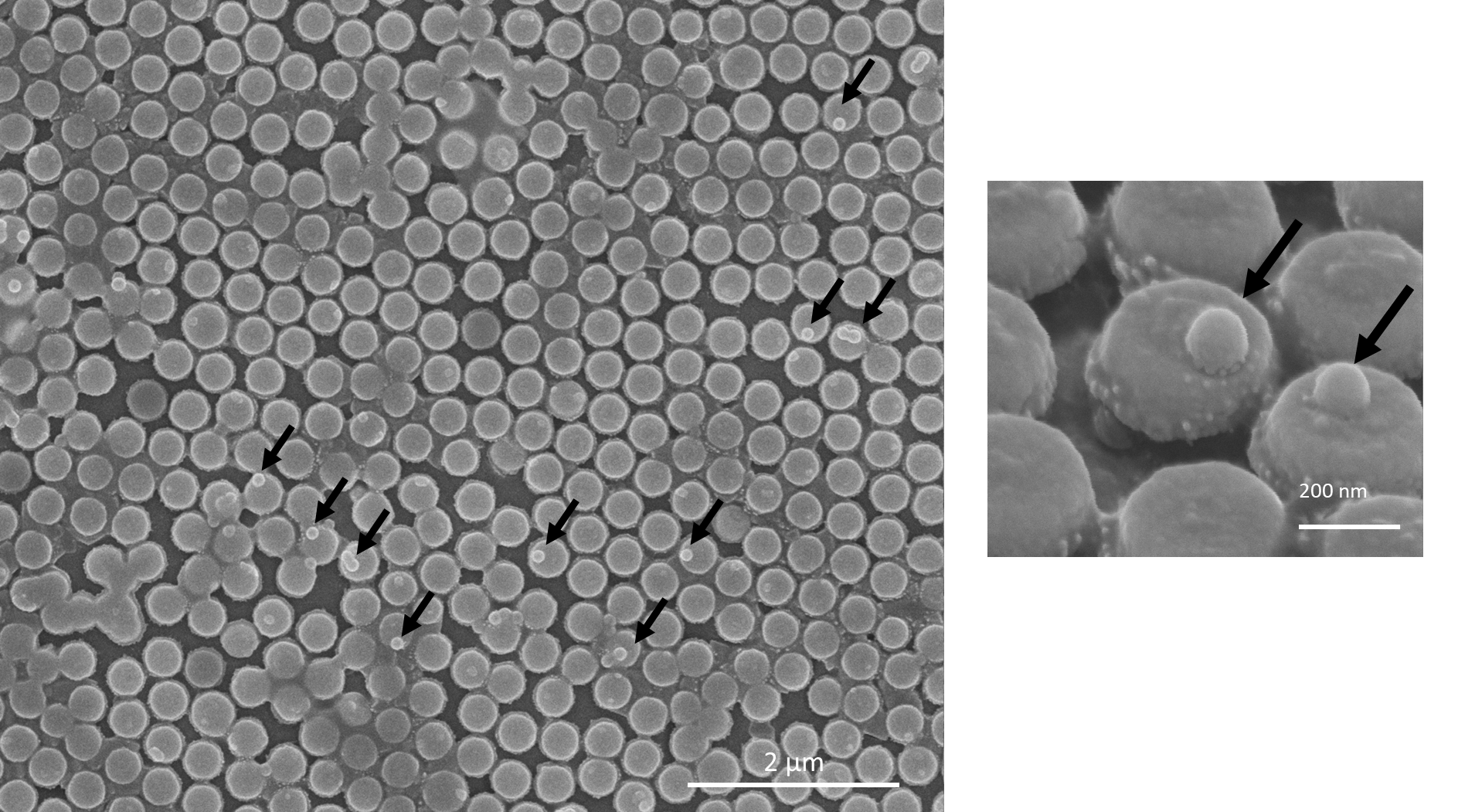

**Supplementary Figure S2.** SEM images of 100 nm PSBs attached on AGNIS.

To determine whether the PANORAMA measurements were performed in the digital single-particle regime, we quantified particle occupancy on AGNIS using randomly selected SEM images of the sensor surface (Supplementary Figure S3). Nanodisks were classified as unoccupied, occupied by one particle, or occupied by multiple particles. Across 13 randomly selected SEM AGNIS regions containing 696 nanodisks, 92.24% of nanodisks were unoccupied, 7.45% contained one particle, and 1.01 % contained two or more particles. This quantitative analysis confirmed that most nanodisks remained unoccupied and that multi-particle occupancy was rare under the assay conditions used in this study.

We further estimated the expected nanodisk occupancy using a Poisson model. From randomly selected SEM images of the sensor regions containing 696 nanodisks (N), approximately 647 were unoccupied ($N_{0})$, 47 (6.75 %) contained one particle and 7 (1.01%) contained 2 or more particles. The mean occupancy per nanodisk was therefore:

$$\lambda=-ln\left( \frac{N_{0}}{N} \right)\approx0.08$$

For a random capture process, the probability of observing $k$particles on a nanodisk is given by:

$$P(k)=\frac{\lambda^{k}e^{-\lambda}}{k!}$$

Using $\lambda=0.08$, the expected probabilities were:

$$P(0)=e^{-\lambda}=0.9224$$

$$P(1)=\lambda e^{-\lambda}=0.0745$$

$$P(\geq2)=1-P(0)-P(1)=0.0031$$

Thus, approximately 92.24% of nanodisks are expected to remain empty, 7.45% are expected to contain one particle, and only 0.31% are expected to contain multiple particles. These results support that the assay operates under sparse occupancy conditions, where PANORAMA-positive events are predominantly isolated single-particle capture events rather than multiple particles occupying the same nanodisk.

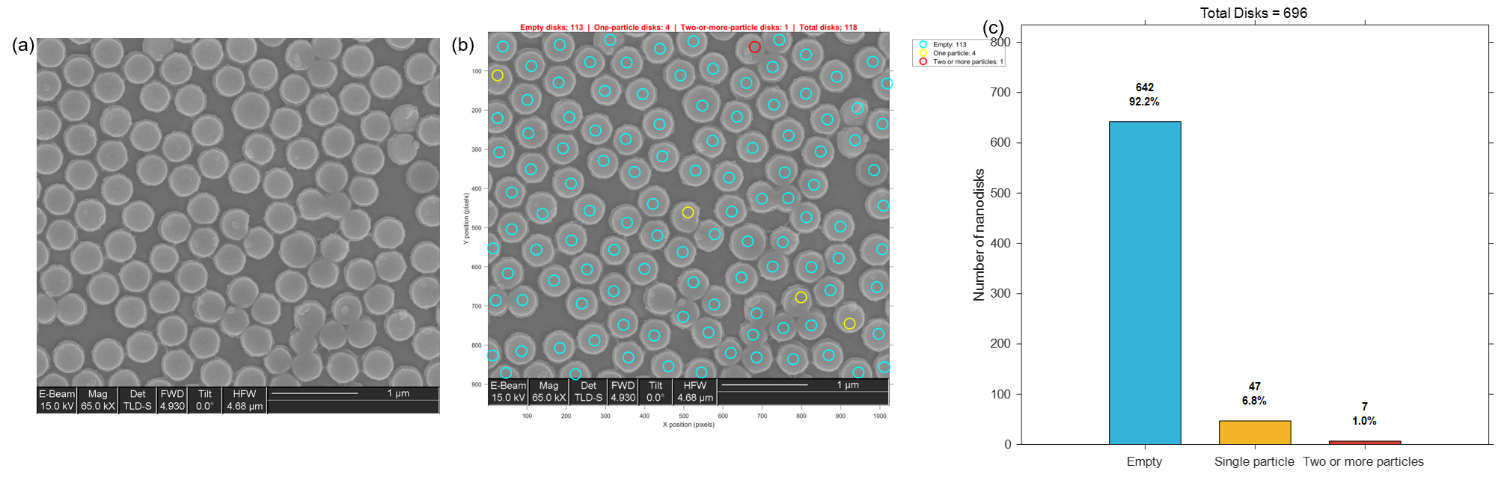

**Supplementary Figure S3.** SEM-based classification of nanoparticle occupancy on nanodisks. (a) Representative SEM image of AGNIS after sensing experiment. (b) Classification of individual nanodisks as empty (cyan), containing a single particle (yellow), or containing two or more particles (red). (c) Combined distribution across all analyzed SEM images. Scale bars: 1 µm.

**Supplementary Note 5**

To directly examine whether the fabrication method influences sensor-level performance, particle detection (25 nm PSBs) was compared using AGNIS substrates fabricated by the petri-dish, LBT, and NSSL methods under identical sample concentration, incubation time, imaging, and analysis conditions. As shown in Supplementary Figure S4, the NSSL-fabricated substrates yielded the highest mean particle count, exceeding the petri-dish method and providing a response comparable to or slightly higher than that of LBT. These results provide direct sensor-level evidence that the improved structural uniformity of sensor fabricated with NSSL results in an improved sensor response. When considered together with the structural and optical uniformity of NSSL, this comparison supports NSSL as a more robust and scalable fabrication approach for AGNIS-based biosensing.

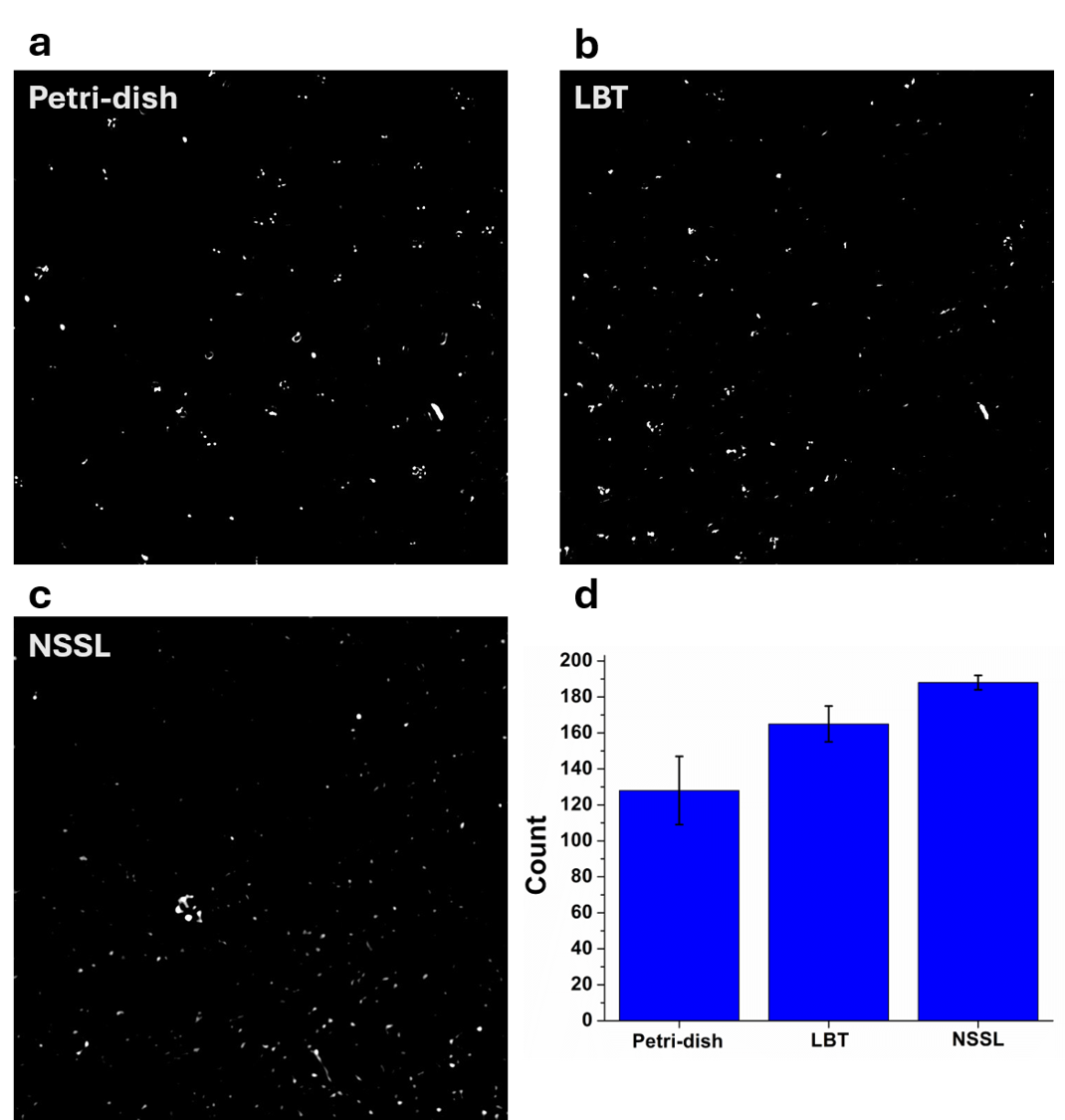

**Supplementary Figure S4.** Comparison of sensing performance of AGNIS substrates fabricated using the petri-dish, LBT, and NSSL methods. (a) Representative PANORAMA images showing the detection of 25 nm polystyrene particles on three sensors fabricated via petri-dish, LBT and NSSL under identical experimental conditions. Scale bars: 10 µm. (b) Mean particle counts obtained from the petri-dish-, LBT-, and NSSL-fabricated substrates. Error bars represent standard deviation from 3 independent measurements.

**Supplementary Note 6:**

The washing protocol was optimized to ensure selective retention of bound sEVs while removing unbound or weakly attached vesicles. Specifically, the AGNIS chips were washed with 1× PBS for 0, 3, 5, 10, and 15 cycles. We observed that unbound sEVs were effectively removed by five wash cycles, beyond which the detected sEV count remained mostly constant (Supplementary Figures S5a–b), indicating stable and specific capture. Supplementary Figure S5c–d shows representative PANORAMA images of detected particles and sEVs before and after washing at the 60 min time point on petri-dish and LBT substrates. SEM imaging was performed following vesicle fixation using a standard protocol^1^. SEM analysis confirmed a one-to-one sEV-to-nanodisk capture configuration, validating specific vesicle binding to individual AGNIS features (Supplementary Figure S5c).

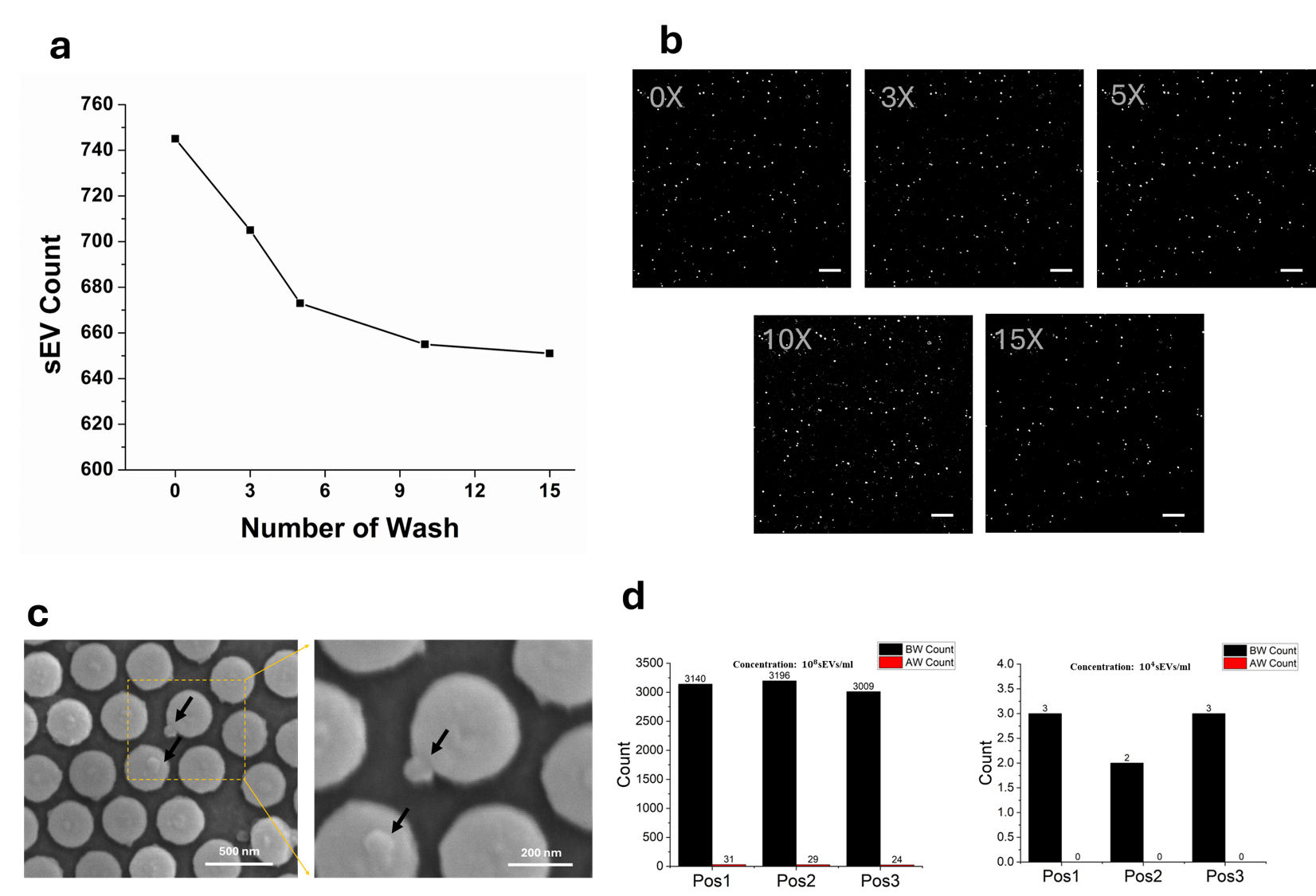

**Supplementary Figure S5. a-b.** Optimization of the washing step PANORAMA of detected sEVs isolated from H460 cancer cell line cultures using **c.** SEM images of sEVs on AGNIS after washing. **d.** Detected sEV counts at highest and lowest concentrations using biotinylated IgG in place of anti-CD9/CD63/CD81 antibodies for surface capture. Scale bar: 20 µm.

**Supplementary Note 7:**

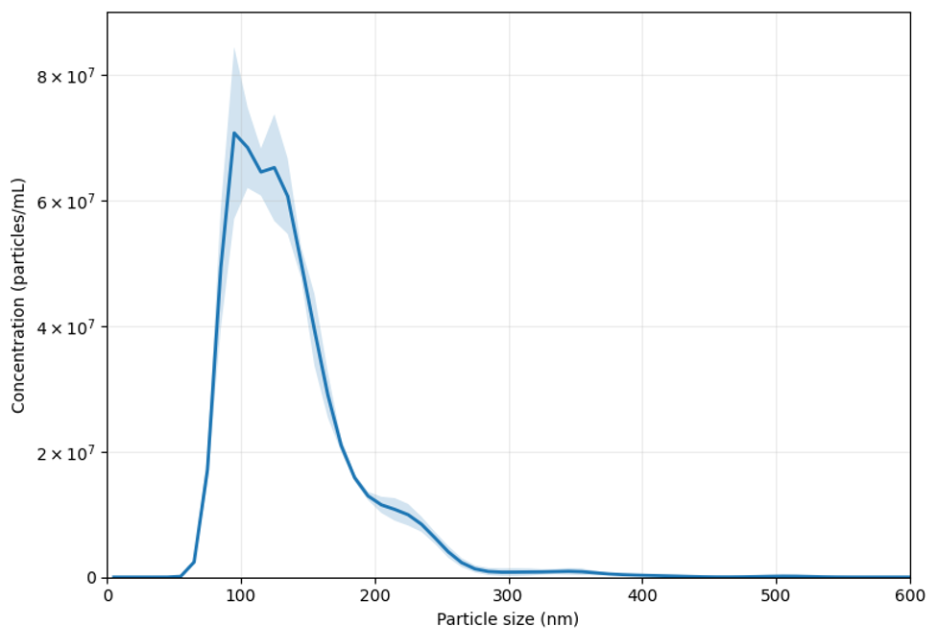

**Supplementary Figure S6.** Nanoparticle tracking analysis (NTA) of the purified sEVs derived from the H-460 cell line. The red trace represents the measured distribution, and the black line represents the smoothed profile.

**Supplementary Note 8**

**Comparison of LOD with representative sEV detection technologies**

To place the analytical sensitivity of the present PANORAMA platform in context, we compared its experimentally determined LOD with representative sEV detection technologies reported in the literature. As summarized in Supplementary Table S1, conventional ELISA and nanoparticle tracking analysis (NTA) typically require approximately 1 × 10⁸ sEVs mL⁻¹, whereas nPLEX has reported an LOD of approximately 4.04 × 10⁵ sEVs mL⁻¹. In comparison, the present PANORAMA platform achieved an LOD of 1 × 10⁴ sEVs mL⁻¹ with a 60 min assay time.

This comparison highlights the high sensitivity of PANORAMA for digital counting of antibody-captured sEVs. However, because each method differs in detection principle, sample preparation, capture strategy, readout format, and LOD definition, the values should be interpreted as approximate order-of-magnitude comparisons rather than direct head-to-head performance comparisons.

**Supplementary Table S1. Comparison of representative sEV detection technologies**

| Technique | LOD | Response Time | Reference |
| --- | --- | --- | --- |
| ELISA | 1×10⁸ sEVs/mL | 2–6 hours | ^2,3^ |
| NTA | ~1×10⁸ sEVs/mL | 10–30 minutes | ^4^ |
| nPLEX | ~4.04×10^5^ sEVs/mL | 1–2 hours | ^3^ |
| PANORAMA | 1×10⁴ sEVs/mL | 60 minutes | ^our work^ |

**Supplementary Note 9:**

**Control Experiments for Validation of miR-21 Detection**

To validate the specificity of the miR-21 detection assay and confirm that the observed fluorescence signals originated from captured sEVs, several control experiments were performed.

**Validation of sEV-specific Fluorescence Detection**

To determine whether the fluorescence signal originated from captured sEVs rather than nonspecific adsorption of plasma proteins or debris, an sEV-depleted plasma sample was prepared by removing vesicles using size-exclusion chromatography [1], which elutes sEVs in the earlier fractions while leaving predominantly soluble plasma proteins and residual debris in the depleted fraction. The sEV-depleted plasma was subsequently incubated with the miR-21 molecular beacon (MB) probe following the same protocol used for whole plasma samples.

Prior to washing, a small number of low-contrast PANORAMA features were observed, with an average PANORAMA contrast of 5.2 ± 1.2%, indicating features substantially smaller than typical sEVs. Following the washing step, only two particles remained within the imaging area, and no detectable fluorescence signal was observed (Supplementary Figure S7a). These results demonstrate that the fluorescence signals observed in whole plasma samples originate predominantly from captured sEVs rather than nonspecific adsorption of plasma proteins or residual debris.

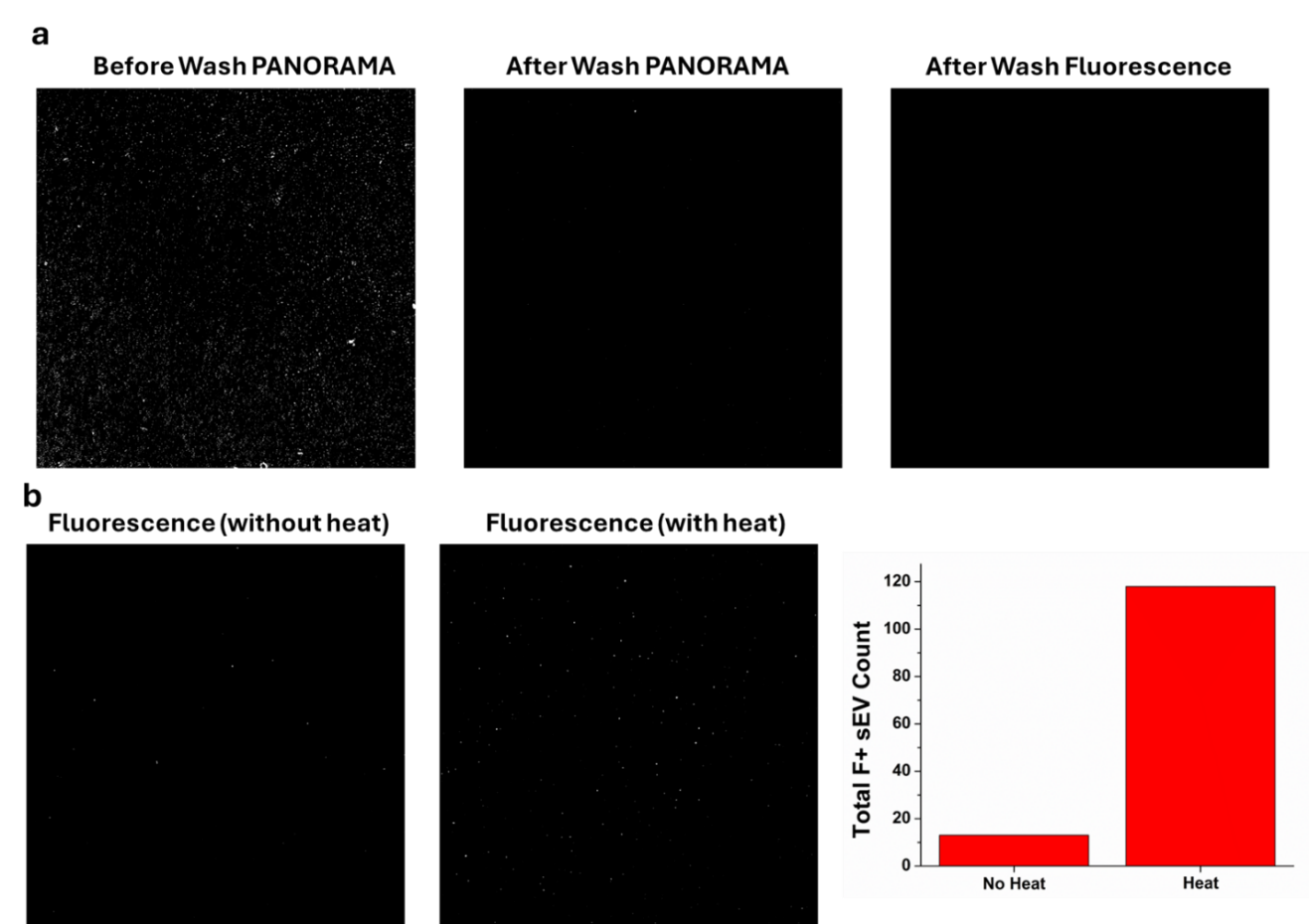

**Supplementary Figure S7:** a. PANORAMA and Fluorescence images of particles from sEV free media. b. Fluorescence images of sEVs and count comparison from liver cancer plasma samples under non-permeabilized and permeabilized conditions.

**Control Experiments for Specificity of PANORAMA-positive Events in Plasma**

Because unpurified plasma contains nanoscale species other than sEVs, including lipoproteins, protein aggregates, nucleoprotein complexes, and other extracellular particles, we performed additional controls to evaluate the specificity of PANORAMA-positive events detected on the sensor surface. PANORAMA detects optical contrast from nanoscale objects bound to the AGNIS surface; therefore, optical contrast alone does not establish vesicle identity. We therefore compared anti-tetraspanin capture surfaces with negative control blocking only, bovine serum albumin (BSA) surface.

Anti-CD9/CD63/CD81-functionalized surfaces produced PANORAMA-positive particle detection from plasma, with 659 ± 23 counts. In contrast, blocking-only surfaces (* BSA only) showed significantly reduced counts of 21 ± 9 (Supplementary Figure S8a). These results indicate that nonspecific adsorption contributes only a minority of the PANORAMA-positive events under the present assay conditions and that anti-tetraspanin capture substantially enriches the detected nanoscale particle population.

To further test whether the detected particles were associated with the sEV-enriched fraction, we analyzed sEV-depleted plasma. Compared with untreated plasma, sEV-depleted plasma showed a marked decrease in PANORAMA-positive events from 659 ± 23 to 2 ± 2 events (Supplementary Figure S8b). This reduction supports the interpretation that most PANORAMA-positive events detected under anti-tetraspanin capture conditions are associated with plasma sEVs or sEV-enriched particles.

These controls support the PANORAMA assay preferentially detects an sEV-enriched nanoscale particle population from plasma rather than nonspecific adsorption alone. However, because plasma contains multiple extracellular nanoparticle classes, we conservatively refer to PANORAMA-only events as “tetraspanin-captured nanoscale particles” unless they are further supported by fluorescence co-localization or additional marker information.

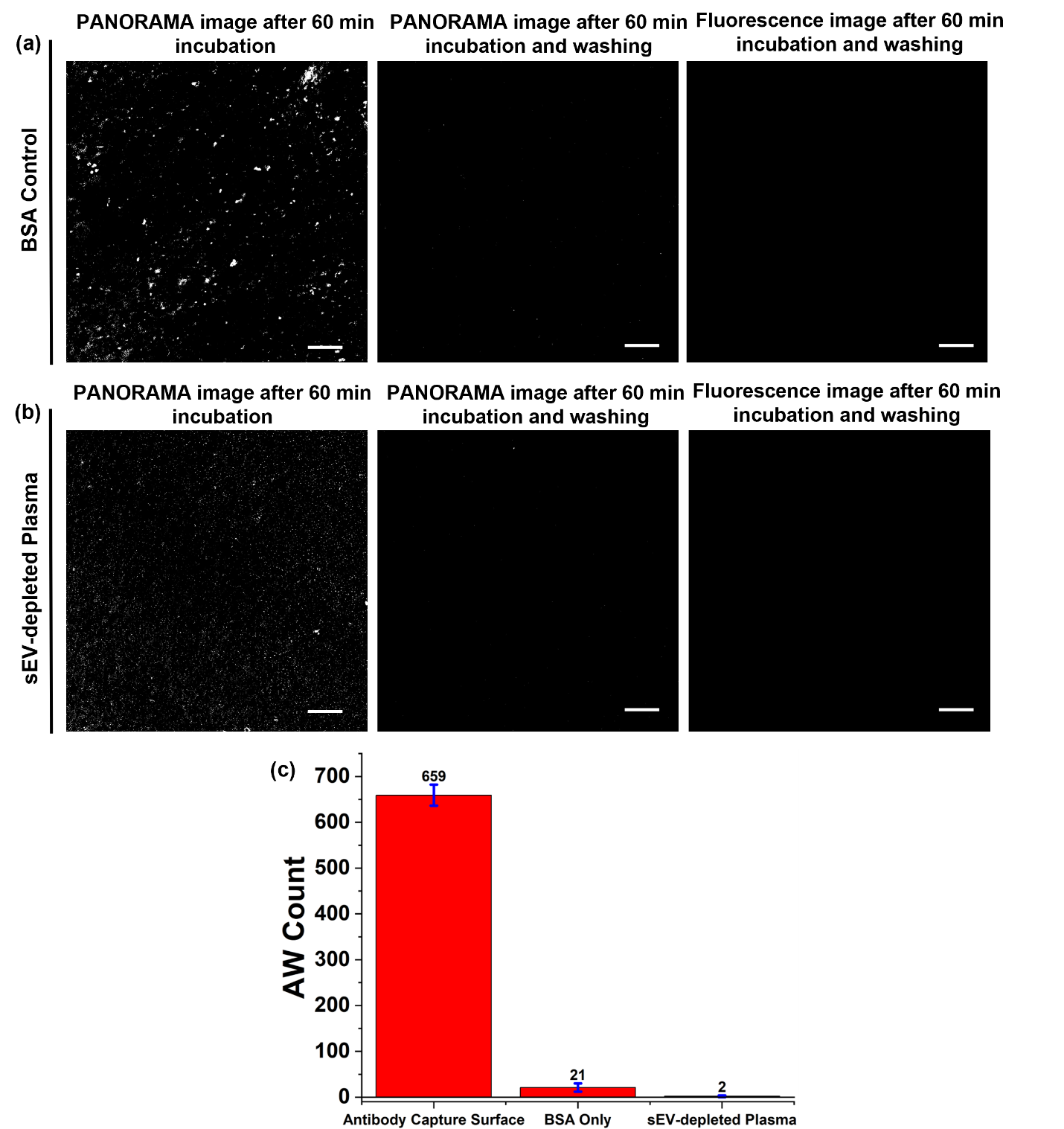

**Supplementary Figure S8:** Specificity controls for PANORAMA-positive nanoscale events in plasma.
PANORAMA-positive event counts were compared across anti-CD9/CD63/CD81 capture surfaces, blocking-only BSA surface, sEV-depleted plasma. Scale bar: 20 µm.

**Effect of Mild Heating on Molecular Beacon Delivery**

A second control experiment was performed to evaluate the effect of the mild heating step used during molecular beacon delivery. Samples incubated with molecular beacons at room temperature were compared with samples incubated at 40 °C under otherwise identical conditions. Heating resulted in an approximately eightfold increase in fluorescence signal (Supplementary Figure S7b), demonstrating that mild heating significantly enhances molecular beacon permeation across the vesicle membrane and improves access to intravesicular miR-21 targets. These results confirm that the heating step is essential for efficient intracellular delivery of the molecular beacon.

**Validation of Fluorescence Specificity Toward Intravesicular miR-21**

To further verify that the detected fluorescence originated from intravesicular miR-21 rather than free miR-21 molecules or plasma components, three additional controls were done: (i) miR-21 molecular beacon solution alone, (ii) plasma alone, and (iii) plasma with miR-21 molecular beacon. Neither the miR-21 molecular beacon solution alone nor the plasma-only sample produced detectable fluorescence signals after the assay. In contrast, the plasma sample with miR-21 molecular beacons generated a clear fluorescence response (Supplementary Figure S9). These results confirm that fluorescence is produced only when the molecular beacon encounters its target miR-21 within the assay environment and further support the specificity of the detection strategy. Also, fluorescence images presented throughout this work were acquired only after the washing step. The washing procedure was intentionally included to remove unbound molecular beacons and weakly adhered vesicles prior to imaging, thereby minimizing background fluorescence and ensuring that the reported fluorescence signals originated from specifically captured sEVs.

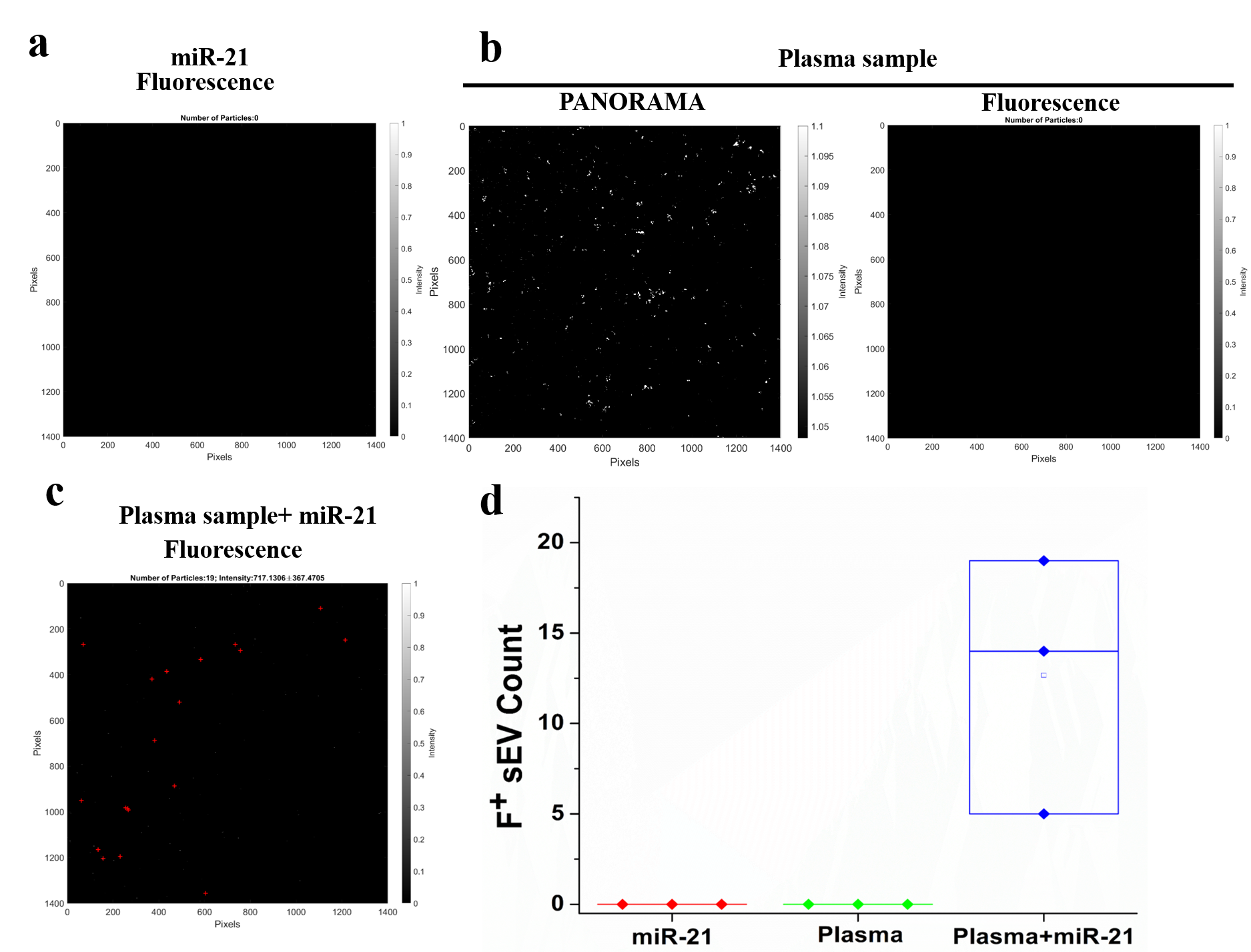

**Supplementary Figure S9:** Validation of the specificity of miR-21 fluorescence detection. a. Fluorescence image of synthetic miR-21 solution alone, showing no detectable fluorescence signal in the absence of plasma and captured sEVs. b. Representative PANORAMA (left) and fluorescence (right) images of plasma alone, demonstrating the absence of detectable fluorescence despite the presence of plasma-derived particles. c. Representative fluorescence image of plasma supplemented with synthetic miR-21, showing a clear fluorescence response following molecular beacon hybridization. d. Quantification of the fluorescence response for the three experimental conditions.
